# SARS-CoV-2 has observably higher propensity to accept uracil as nucleotide substitution: Prevalence of amino acid substitutions and their predicted functional implications in circulating SARS-CoV-2 in India up to July, 2020

**DOI:** 10.1101/2020.10.07.329771

**Authors:** Subrata Roy, Himadri Nath, Abinash Mallick, Subhajit Biswas

## Abstract

SARS-CoV-2 has emerged as pandemic all over the world since late 2019. In this study, we investigated the diversity of the virus in the context of SARS-CoV-2 spread in India. Full-length SARS-CoV-2 genome sequences of the circulating viruses from all over India were collected from GISAID, an open data repository, until 25^th^July, 2020. We have focused on the non-synonymous changes across the genome that resulted in amino acid substitutions. Analysis of the genomic signatures of the non-synonymous mutations demonstrated a strong association between the time of sample collection and the accumulation of genetic diversity. Most of these isolates from India belonged to the A2a clade (63.4%) which has overcome the selective pressure and is spreading rapidly across several continents. Interestingly a new clade I/A3i has emerged as the second-highest prevalent type among the Indian isolates, comprising 25.5% of the Indian sequences. Emergence of new mutations in the S protein was observed. Major SARS-CoV-2 clades in India have defining mutations in the RdRp. Maximum accumulation of mutations was observed in ORF1a.

Other than the clade-defining mutations, few representative non-synonymous mutations were checked against the available crystal structures of the SARS-CoV-2 proteins in the DynaMut server to assess their thermodynamic stability. We have observed that SARS-CoV-2 genomes contain more uracil than any other nucleotide. Furthermore, substitution of nucleotides to uracil was highest among the non-synonymous mutations observed. The A+U content in SARS-CoV-2 genome is much higher compared to other RNA viruses, suggesting that the virus RdRp has a propensity towards uracil incorporation in the genome. This implies that thymidine analogues may have a better chance to competitively inhibit SARS-CoV-2 RNA replication than other nucleotide analogues.

## 1. Introduction

The world is in a pandemic situation due to an outbreak of highly infectious human to human transmissible virus, named SARS-CoV-2. Since the first novel pneumonia case in Wuhan, China 17,396,943 confirmed cases with 675,060 deaths were reported until 1^st^ August 2020 (WHO, 2020). The virus was found to be a strain of beta-coronavirus and related to SARS-like BAT coronaviruses, bat-SL-CoVZC45 and bat-SL-CoVZXC21 with 88% similarity; 79.5% homology with SARS, and 50% with MERS (Lu *et al.*, 2020; Wu *et al.*, 2020). The virus originated from its root (Wuhan) and is changing while spreading throughout the world. It is an RNA virus with a higher mutation rate over DNA viruses. Therefore, characterisation of circulating strains is important to correlate with disease pathogenesis and outcome; decide on treatment strategies as well as to obtain real-time background information for developing effective vaccines and antivirals.

Over the few months, several studies have been published which reported some novel mutations in Indian isolates. In this study, whole-genome mutation analysis has been done for a total of 1878 sequences that have been reported from India. Nucleotide changes that have introduced non-synonymous changes in the gene have been considered for analysis. Here, all the sequences were analyzed to find in which clades they fit best in respect to the global scenario. In this study, we have presented a clear understanding of the mutations prevalent in SARS-CoV-2 isolates from India till the end of July 2020.

## 2. Materials and methods

Since the outbreak of SARS-CoV-2 in China in December 2019, GISAID (https://www.gisaid.org/) has become a global repository for coronavirus genome sequences. For this study, 365 full genome sequences from India until 25^th^ May, 2020 were collected from the GISAID server. Later 1513 more sequences were added to this study to compare the pattern of SARS-CoV-2 spread across India. Sequences were aligned using the MEGA X software (Kumar *et al.*, 2018). Position of amino acids was defined with respect of the first viral genome sequence, named ncov2019-Wuhan-hu-1/2019 (GenBank accession no: MN908947), taken as the parental or reference strain in the multiple sequence alignment. Analysis of data was done using Bioedit (Hall, Biosciences and Carlsbad, 2011).

The functional implications of non-synonymous mutations were predicted in the DynaMut (Rodrigues *et al.*, 2018) server utilizing the available crystal structure data specific to SARS-CoV-2 proteins. Several mutations which were found to be selective in the population; reported previously as important, or used as a marker to define different clades, were studied in DynaMut to study their functional importance. DynaMut score ΔΔG for each mutation was considered to predict whether the mutation is stabilizing or not. In order to predict the flexibility of a protein with a given mutation, the free entropy change was considered and this also gives the prediction of future selectivity of the said mutation. Flexibility and rigidity are the key contributors to protein function. Consequently, in higher temperature fluctuations, a rigid protein structure is beneficial for protein structure stability rather than a flexible structure (M, 1987). In our DynaMut studies, ΔΔSVib ENCoM is the change in vibrational entropy energy between wild-type and mutant protein. The value of ΔΔSVib ENCoM predicted in the DynaMut server for each point mutation signifies the change in the molecular flexibility of the protein. Negative value indicates a decrease in flexibility and vice versa. This means mutations that confer potential structural rigidity to the proteins (ΔΔG value positive; ΔΔSVib ENCoM, negative) might compensate for higher temperature oscillations. Hence based on the calculative predictions, such mutations may constitute a stable conformation of the proteins in the virus evolution.

In this study, the potential impact of a point mutation in the protein structure was also predicted via free energy-based (ΔG) calculative method in DynaMut server. Low free energy value signifies a stable protein conformation and high free energy value for unstable protein conformation. So, if an amino acid substitution lowers the free energy value from the wild type, the mutation dictates a stabilizing conformation of the protein. From the DynaMut server, we obtained a ΔΔG value which means a difference between ΔG wildtype and ΔG mutation (ΔΔG= ΔG wildtype-ΔG mutation).

## 3. Results

### 3.1. Defining types of SARS-CoV-2

Overall, a high level of sequence identity throughout the 29 kb genome was observed considering the single nucleotide polymorphisms that resulted in change of amino acids in comparison to the reference sequence (MN908947.3). Other than the clade-defining ones, mutations that occurred on more than ten occasions (considering all sequences), have been used in our analysis. Others have been ignored as they may be sequencing artefacts. According to previous reports, the earliest sequences of SARS-CoV-2 from China belonged to clade O. In addition to the ancestral type (O), there were 10 derived types. Among them five derived types had high frequencies in India, namely O, B, B1, A1a, and A2a up to early May, 2020 (Biswas and Majumder, 2020).

In total, 1190 isolates contained both D614G (nt 23403 A>G) in spike protein along with P323L (nt 14408C>T) in RdRp of ORF1b which group them under the A2a clade. The viruses of clade A2a constituted 63.4% of the total reported sequences in India. Twenty-seven isolates were A3 type (1.4%), containing the clade-defining mutations V378I (nt 1397G>A) and L3606F (nt 11083G>T) in ORF1a. Total 82 isolates from India contained L84S (nt 28144T>C) mutation in ORF8 which is a clade-defining mutation for B type. Among them, 73 isolates also had S202N (nt 28878G>A) mutation in N gene as well. These 73 isolates belonged to B4 type (3.9%). Among the rest 9 isolates, 8 isolates were of B type (0.4%) (Fig: 1). Therefore, it appears that B4 emerged from B and predominated among the B types over the duration of our study. The other isolate with the L84S mutation belonged to A2a type (Supplementary Table 1).

**Fig 1:**
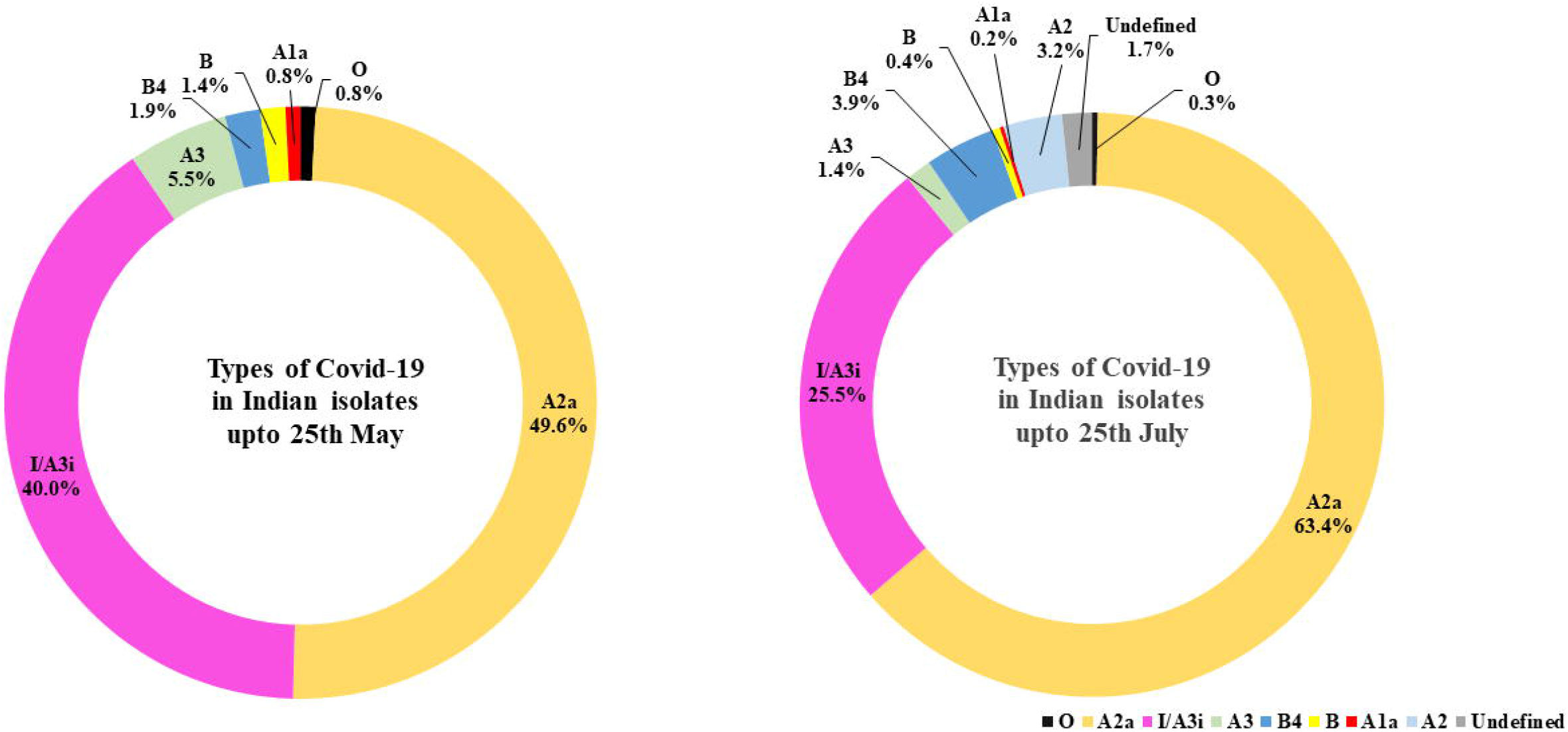
Distribution of different types of SARS-CoV-2 among the Indian population. **(a)** Most of the isolates belong to 2 major types of virus, A2a and I/A3i. A new cluster of viruses I/A3i was found to be getting fixed in the population until 25^th^ May 2020. **(b)** The extended study revealed that the percentage of I/A3i decreased from 40% to 25% by 25^th^ July 2020 and A2a became the predominant type.

Up to 1^st^ May, 2020 only 8 out of the total 56 Indian isolates reported till that time, contained some novel mutations i.e. P13L (nt 28311C>T, N gene), A97V (nt 13730C>T in RdRp ORF1b) and T2016K (nt 6312C>A) along with L3606F (nt 11083G>T) in ORF1a. Later, with more and more sequences being available, a different cluster gradually emerged with the above three clade-defining mutations. This clade had been named I/A3i (Jolly *et al.*, 2020). In case of few isolates, any one of the above-mentioned mutations could not be determined but rest were present, so those genomes had been included within the I/A3i type. Interestingly, this cluster was found to constitute 40% of the total reported sequences in India until 25^th^ May. However, the scenario changed on extending our study to the end of July. The I/A3i type decreased to 25.5% of the total sequence observed, while A2a subtype increased from 50% to 63% of the total study population. Emergence of another subtype was observed in the form of A2 in case of the SARS-CoV-2-infected Indian population. Until the end of May, no A2 subtype was observed which carried only the D614G mutation in the spike protein. But extended study identified 61 sequences (3.2%) of A2 subtype in the Indian population. In case of 31 Indian isolates, specific clade couldn’t be assigned due to lack of sufficient or confirmed sequence information in the clade-defining positions.

### 3.2. Other non-synonymous mutations

Our study has identified some other mutations among the different clades which may play an important role in the course of viral genome divergence (Table 1). For instance, besides the clade-defining mutations, two other mutations were observed in high frequency (n=20/27 each) in the A3 cluster; i.e. R207C and M2796I in nsp2 and nsp4 respectively.

**Table 1:**
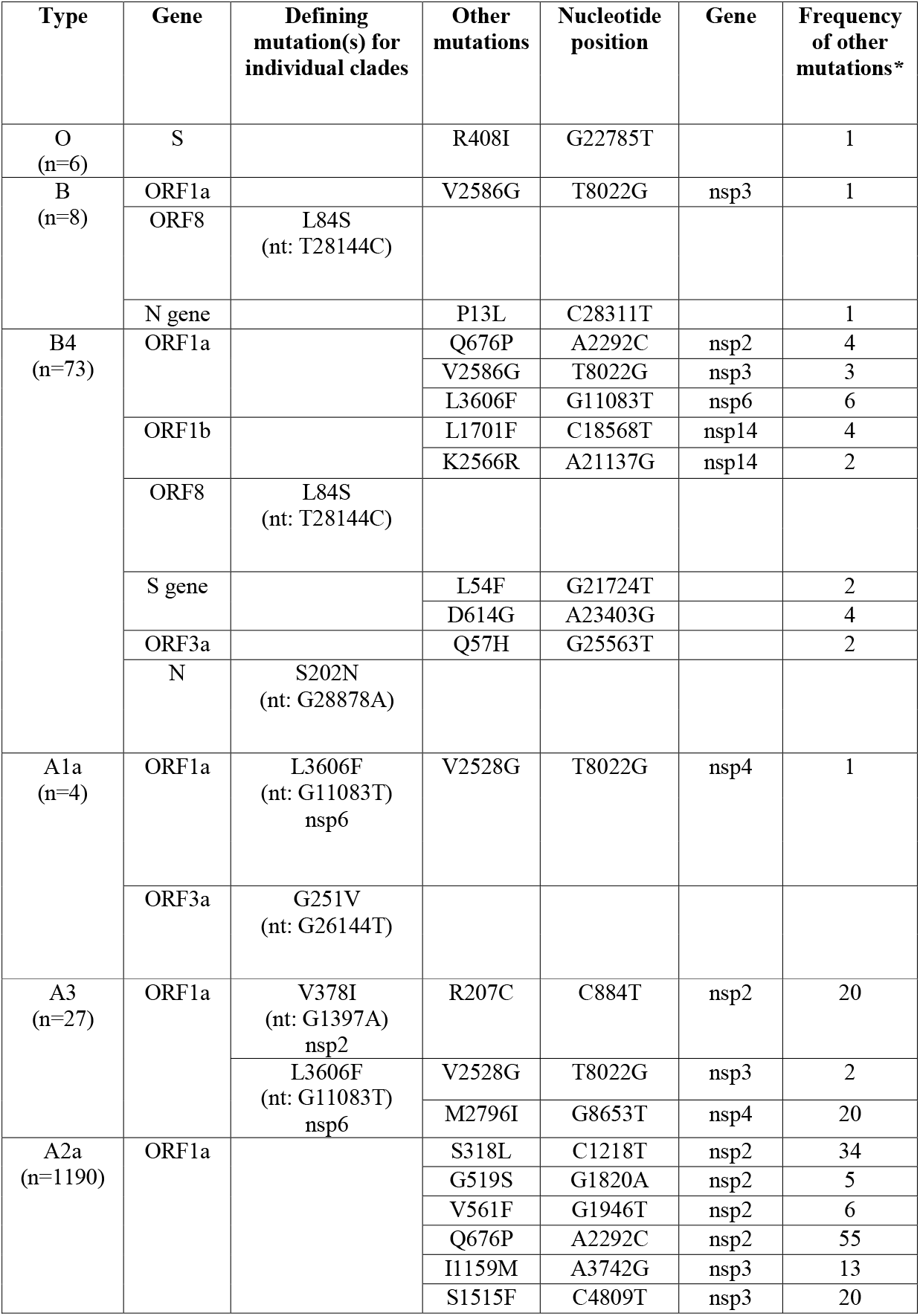
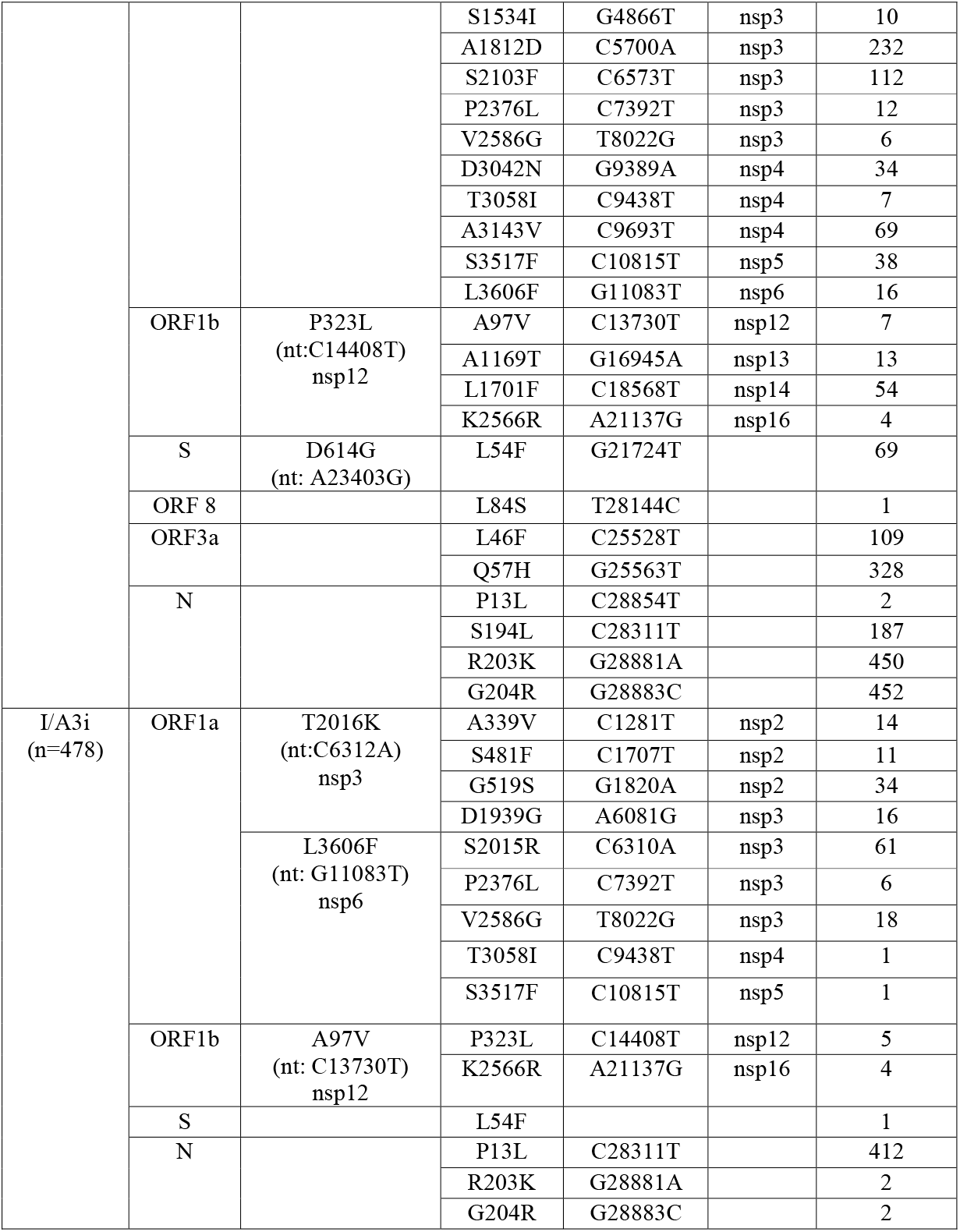
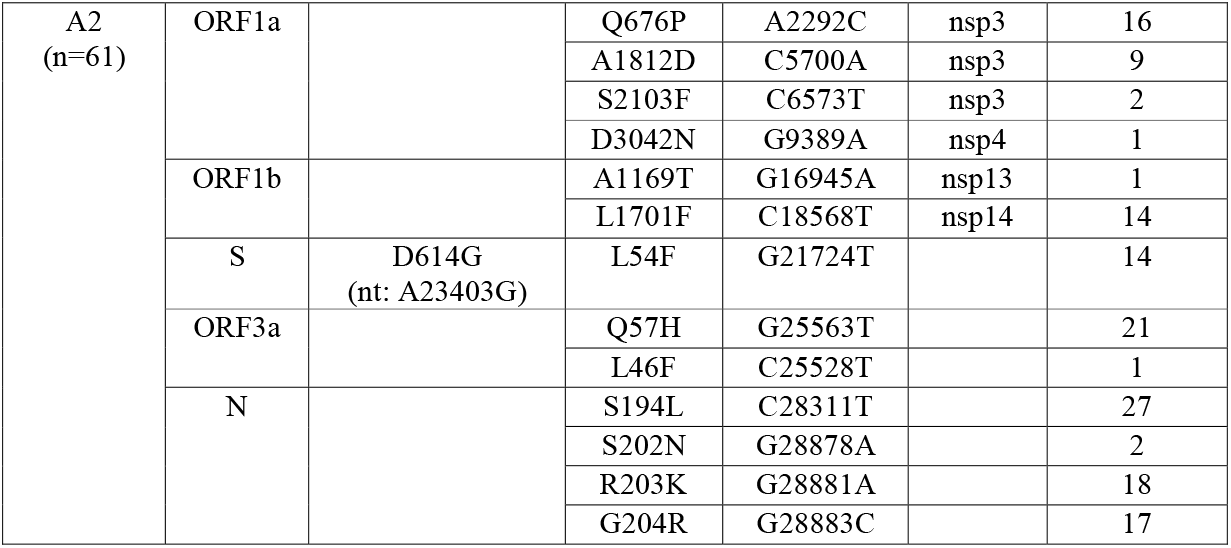
Non-synonymous mutations and corresponding frequencies across the different clades of Indian SARS-CoV-2 isolates. *Except the R408I mutation (n=1) in O type S protein, non-synonymous mutations with cumulative frequency of ≥10, have only been considered.

Two consecutive amino acid changes R203K and G204R could be observed in the nucleocapsid phosphoprotein (N protein) of approx. 30% A2 and 37% A2a Indian isolates. These mutations were previously reported to be also abundant in the USA (Joshi and Paul, 2020). Over the course of infection A2a type has acquired highest number of mutations in ORF1a such as, S318L (n=34/1190) and Q676P (n=55/1190) in nsp2; A1812D (n=232/1190) and S2103F (n=112/1190) in nsp3; D3042N (n=34/1190) and A3143V (n=69/1190) in nsp4 and S3517F (n=38/1190) in nsp5 of ORF1a (Fig:2). Similarly, A2a type also showed non-synonymous changes in other regions, like Q57H (n=328/1190) and L46F (n=109/1190) in ORF3a and S194L (n=187/1190) in the N gene. Another mutation in N gene P13L was observed abundantly in I/A3i type (n=412/478). Other mutations in ORF1a, like G519S (n=34/478) and S2015R (n=61/478) were observed with a high frequency of occurrence in I/A3i clade. The newly emerged A2 type in India also showed high number of mutations in ORF1a as observed in case of the A2a type (details in Table 1).

**Fig 2:**
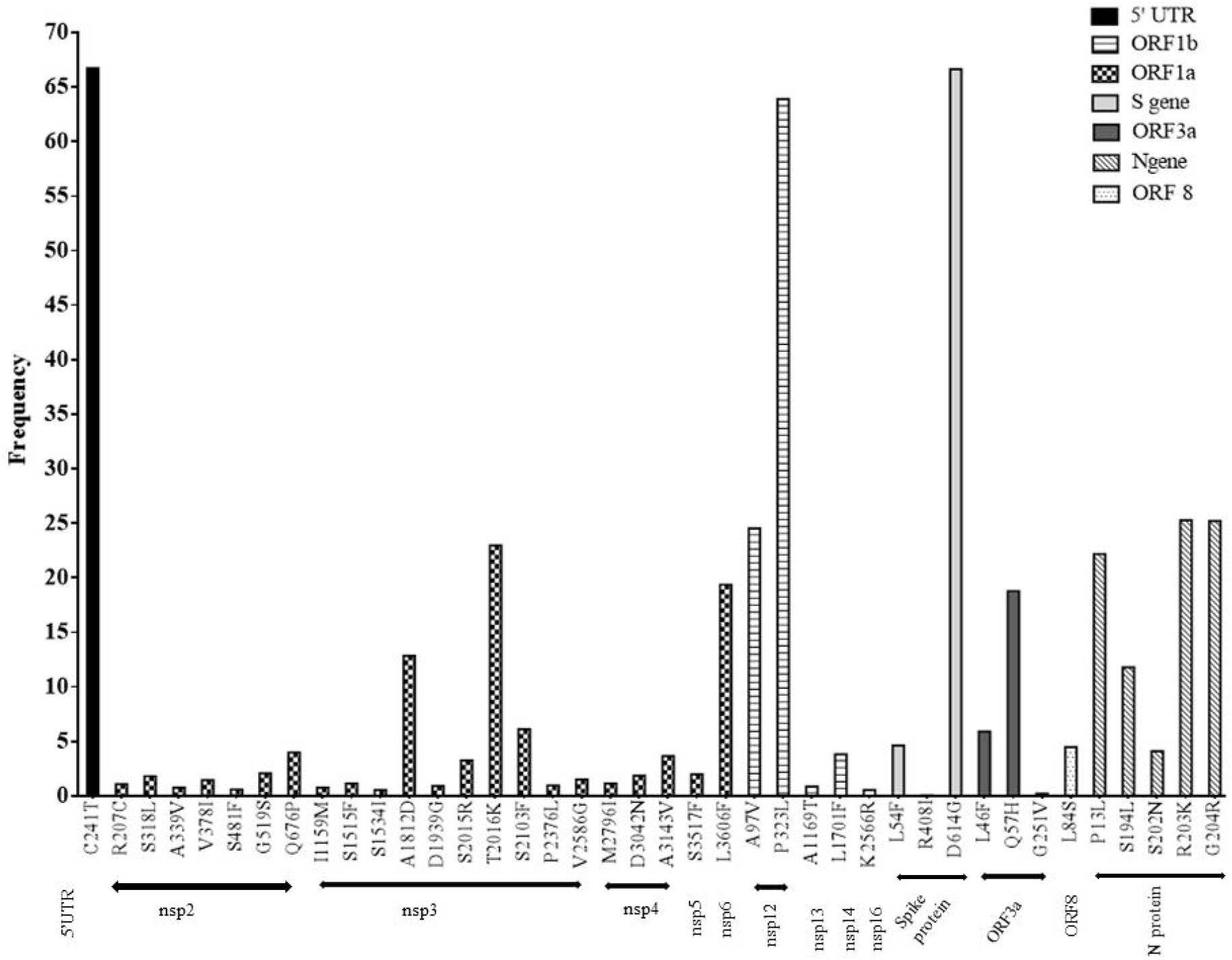
Distribution of non-synonymous mutations across the Indian SARS-CoV-2 genomes. Highest accumulation of mutation can be observed in ORF1a compared to the overall genome. Among the non-structural proteins, nsp3 tends to accumulate the greatest number of mutations.

### 3.3. DynaMut Analysis

DynaMut analysis was done to predict the effect of a point mutation on respective protein stability and molecular flexibility. In the case of spike protein, the mutation D614G had a structurally stabilizing effect on the protein in terms of free energy change (ΔΔG) (Table 2) and is a clade-defining stable mutation for A2 and A2a types. Similarly in case of nsp12 (RdRp), two clade-defining mutations A97V and P323L also appeared to have stabilizing effect on the RNA polymerase, with decreased molecular flexibility (Karshikoff, Nilsson and Ladenstein, 2015). A representative mutation in nsp3 i.e. I1159M was found to be destabilizing and occurred at lower frequency(n=14). Interestingly, DynaMut analysis revealed R408I mutation in S protein to be a stabilizing one but this mutation appeared on a single occasion in a O type virus among all sequences analyzed (Table 2).

**Table 2:** Analysis of the mutations and their stability by DynaMut. *Frequency of each mutation was calculated including the 31 Indian sequences which were undefined and could not be assigned to any particular clade.

|  | **Mutation** | **DynaMut $\Delta\Delta G$**<br>(kcal/mol) | **$\Delta\Delta S_{Vib}$ ENCoM ($\Delta$ Vibrational**<br>**Entropy Energy between**<br>**Wild-type and Mutant)**<br>(kcal.mol<sup>-1</sup>. K<sup>-1</sup>) |
| --- | --- | --- | --- |
| S gene<br>(PDB ID:<br>6VSB) | L54F<br>(n=87)* | 1.746<br>(Stabilizing) | -4.764<br>(Decrease of molecule flexibility) |
|  | R408I<br>(n=1) | 1.857<br>(Stabilizing) | -4.408<br>(Decrease of molecule flexibility) |
|  | D614G<br>(n=1256) | 1.128<br>(Stabilizing) | -4.531<br>(Decrease of molecule flexibility) |
| ORF 1a | I1159M<br>nsp3<br>(PDB ID: 6WEY)<br>(n=14) | -0.258<br>(Destabilizing) | 0.309<br>(Increase of molecule flexibility) |
|  | S3517F<br>nsp5<br>(PDB ID: 6Y84)<br>(n=39) | 1.041<br>(Stabilizing) | -0.249<br>(Decrease of molecule flexibility) |
| ORF1b | A97V<br>nsp12<br>(PDB ID: 6M71)<br>(n=462) | 1.397<br>(Stabilizing) | -5.146<br>(Decrease of molecule flexibility) |
|  | P323L<br>nsp12<br>(PDB ID: 6M71)<br>(n=1210) | 1.540<br>(Stabilizing) | -4.820 kcal.mol<sup>-1</sup>. K<sup>-1</sup><br>(Decrease of molecule flexibility) |

### 3.4. The propensity of the nucleotide changes

The SARS-CoV-2 viral genome is made up of 62% A+U content while the G+C content is 38%. In our study, we have observed that uracil content (32% of the total genome) was much higher in SARS-CoV-2 compared to other RNA viruses like Dengue (20%) or Chikungunya (20%) (Fig: 3). Furthermore, it was found that majority of the non-synonymous mutations occurred due to change of other nucleotides to uracil (64% of the total non-synonymous mutations analysed). The highest rate of substitution was observed as cytosine to uracil (40% of the total number of mutations). (Fig: 4)

**Fig 3:**
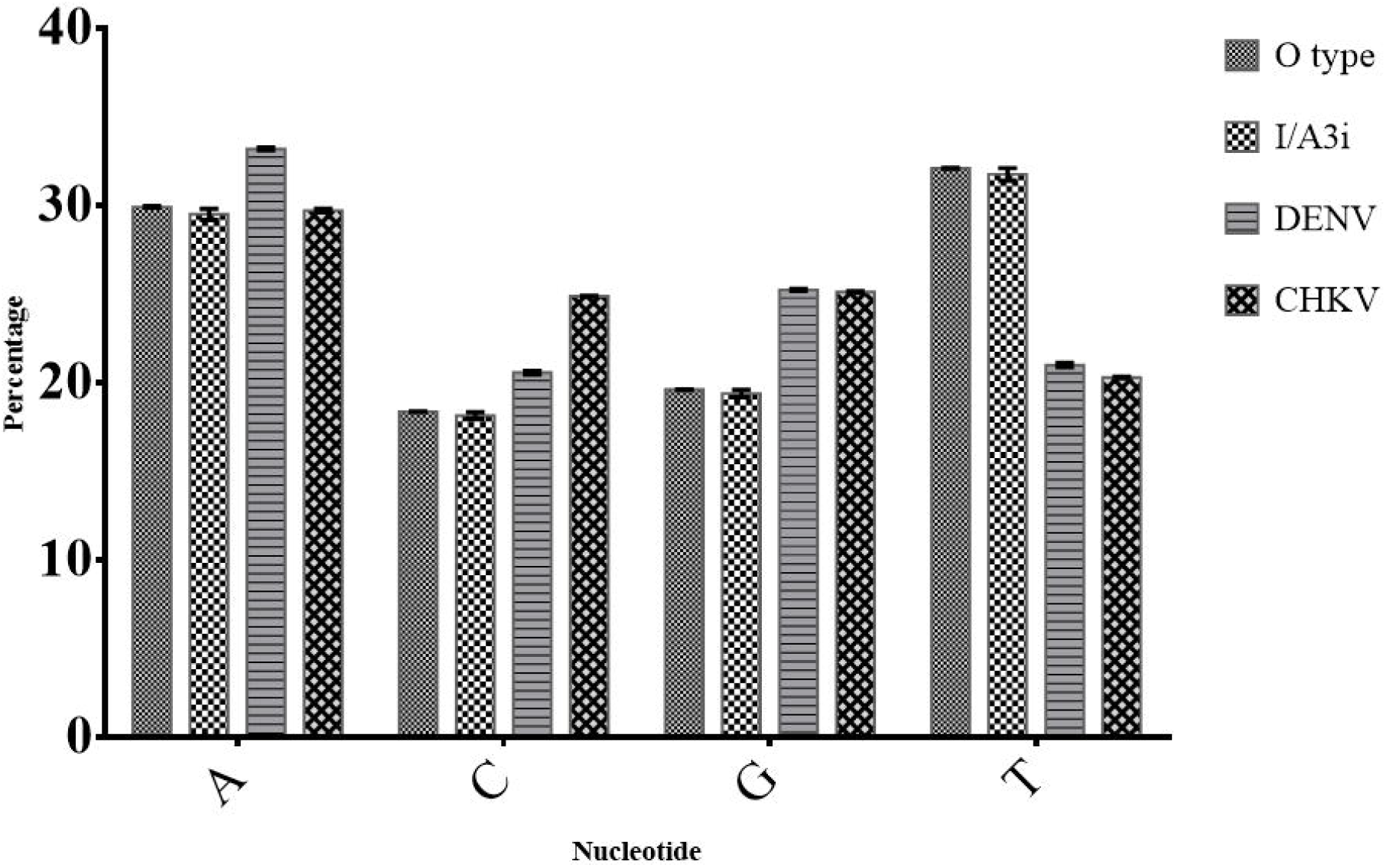
Comparison of nucleotide composition of SARS-CoV-2 RNA backbone with that of two other prevalent RNA viruses in India. The frequency of uracil is highest among all four nucleotides in SARS-CoV-2 genomes. This holds true for both older and recently emergent types of SARS-CoV-2 Indian sequences. Average nucleotide distribution in the RNA backbone of each virus was calculated from three sequences for each virus. Error bars represent SD among the three sequences of each virus used in the comparison.

**Fig 4:**
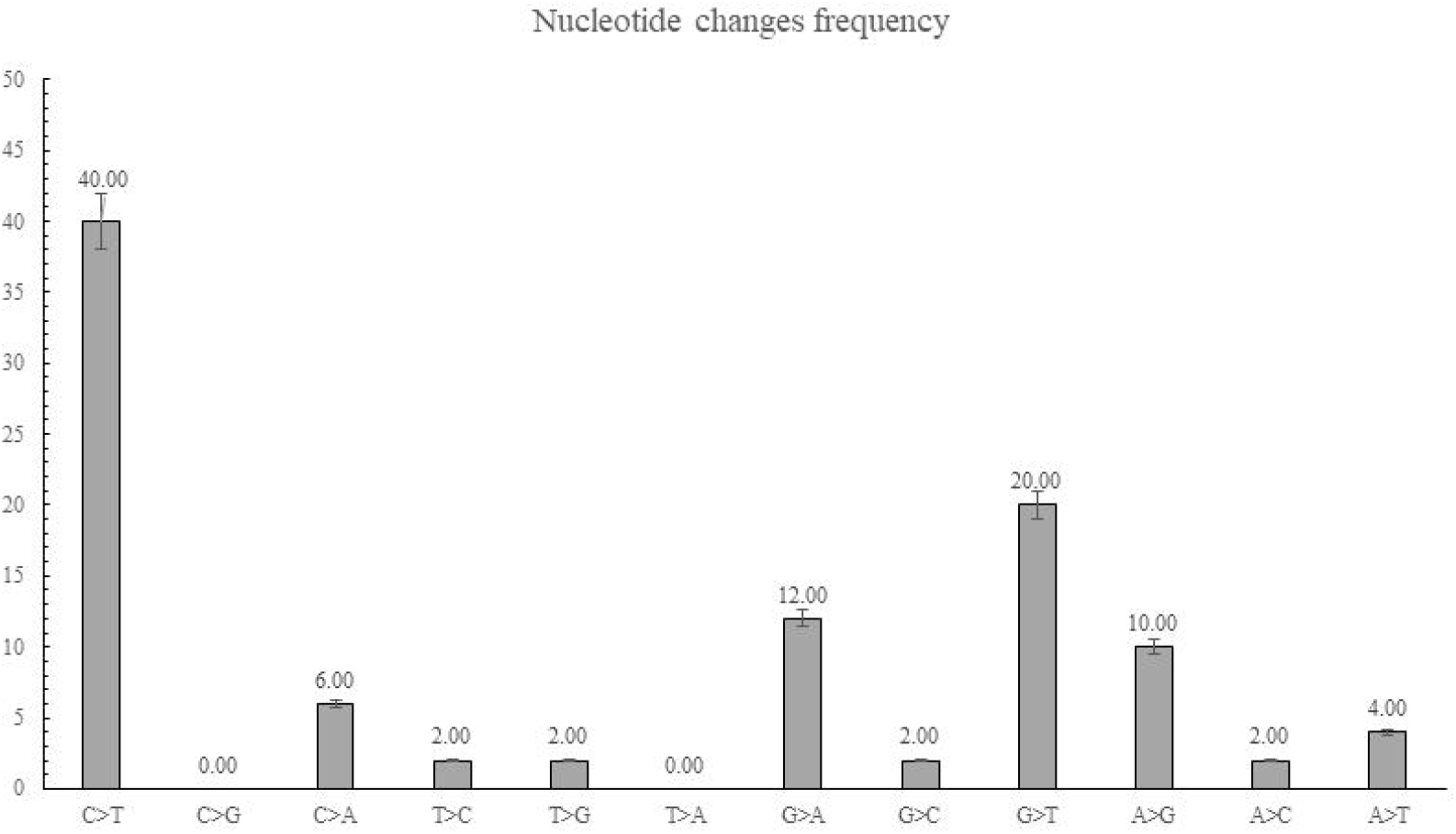
Frequency of non-synonymous nucleotide substitutions expressed as a percentage of the mutations resulting in amino acid substitutions. Most frequent changes in nucleotides were observed in form of C>T (40%). Substitution of G to T was recorded second-highest, sharing 20% of the total non-synonymous mutations. Overall, 64% of all the non-synonymous mutations were substitutions to uracil/thymidine.

## 4. Discussion

Since the outbreak of the SARS-CoV-2, it spread rapidly to more than 200 countries in different continents. Notably, the USA and Western European countries like Spain and Italy had seen a high rate of mortality. Until August 1^st^, 2020 India has reported 1,695,988 confirmed cases and 36,511 deaths due to COVID-19 (WHO, 2020). Compared to the global scenario where death rate due to SARS-CoV-2 was 3.8%, a densely populated country like India had reported only 2.15% mortality due to this highly transmissible virus.

This study was done to understand whether the observed geographical variations in the prevalence of infection, had any relation with particular SARS-CoV-2 clusters. The study was done to assess pathogen evolution with disease transmission. Studies have revealed that a particular subtype A2a, had spread rapidly throughout the European and North American continents and entered East Asia in January 2020. The spread of this subtype rapidly increased from 2% to 60% within 10 weeks (Bhattacharyya *et al.*, 2020).

Most of the sequences from India belonged to the A2a subtype before 1^st^ May. But another cluster of sequences was reported later and classified as I/A3i subtype (Jolly *et al.*, 2020). Surprisingly, the spread of the I/A3i subtype escalated from 14% to 40% of the total reported sequence within 3 weeks (1^st^ May to 25^th^ May 2020). Later the spread of I/A3i decreased to 25.5% of the total infections by the end of July. Over this time, the A2 type (3%) of SARS-CoV-2 emerged in the Indian population, which contained only D614G mutation in the spike protein as the clade-defining mutation. In the time period covered, mainly two types of SARS-CoV-2 isolates were prevalent in India i.e. A2a (63.4%) and I/A3i (25.5%). Other isolates mainly belonged to B4 (3.9%), A3 (1.4%), B (0.4%), O (0.3%) and A1a (0.2%) types (Fig 1).

The most-mentioned mutation in the spike protein, D614G, was observed in all the Indian A2 and A2a sequences (1256 out of 1878 genomes). It has been suggested as one of the major factors behind the virulence of the virus (Korber *et al.*, 2020). However, the position of this mutation is far away from the RBD. Reports suggested that D614G mutation at the junction of the S1 and S2 subunits of S gene introduces an additional cleavage site in the S protein (Bhattacharyya *et al.*, 2020). It has been predicted to reduce host immune response by producing “decoy” fragments that bind to and inactivate antiviral antibodies (Park *et al.*, 2016). This was anticipated to help the virus evade the primary immune response and establish an infection rapidly. It had been experimentally shown that D614G mutation can also increase infectivity substantially by facilitating receptor-ligand interactions (Zhang *et al.*, 2020). Two other notable mutations P323L in RdRp and C241T (synonymous nucleotide change) in 5’ UTR are co-evolving with this mutation. Coronaviruses contain sub-genomic identical 5’leader sequence which plays a role in virus replication. It will be interesting to see whether these changes have any influence on altering the efficiency of viral replication or not.

Two highly predominant clades I/A3i and A2a in India contain distinct mutations in nsp12, i.e. A97V and P323L respectively. These non-synonymous mutations have been used to define clade as well. So, the virus is possibly adapting through gain of mutations in the RdRp and possibly towards more effective replication potential. This proposition also needs experimental validation.

In our analysis, 31 isolates could not be assigned to any specific clade as they either contained mutations overlapping different clades or ‘N’/s (i.e. nucleotide/s could not be determined by sequencing) at clade-defining areas of the genomes. One such isolate from Gujrat (hCoV-19/India/GBRC24b/2020|EPI_ISL_437454|2020-04-26) showed a unique combination of mutations from two different types. This isolate contains P323L (nt 14408C>T) in nsp12 but not D614G (nt 23403A>G) in the S gene which is required to define it as an A2a type. On the other hand, this also contains T2016K (nt 6312C>A) in nsp3 and A97V (nt 13730C>T) in nsp12 but does not have L3606F (nt 11083G>T) in nsp6. So, it could not be established as a genuine I/A3i type also. It appears to be either a hybrid/recombinant of the two types. Alternatively, the patient might have been infected with two different types of SARS-CoV-2 and this peculiar genome sequence is the artefact of sequence assembly of reads generated from mixed sequences.

As the prevalence of A2a and I/A3i is increasing rapidly, we need to observe closely for these kinds of isolates. It has been observed that of P13L (nt C28311T) and S194L (nt C28863T) mutations in nucleocapsid protein are emerging at high frequency among recently uploaded sequences from India. For instance, the distribution of the P13L mutation in I/A3i clade (n=412 out of 478) suggests that it is perhaps evolving towards becoming a clade-defining mutation for I/A3i. S194L mutation was only observed among A2a and A2 types of the Indian isolates.

Non-synonymous mutations that were encountered on ≥10 occasions were considered in our study. DynaMut analysis was performed for those SARS-CoV-2 proteins for which the crystal structure data were available. Mutations such as D614G in S protein and A97V and P323L in nsp12 were found to be stabilizing by the DynaMut analysis. Their predicted stability was further supported by the observed high frequency of these mutations suggesting that these mutations are getting fixed in the population. Interestingly, the R408I mutation (nt G22785T, n=1) in S protein was predicted as a stable mutation by the DynaMut programme and had been previously reported as a potential RBD-altering mutation (Saha *et al.*, 2020). However, this mutation did not appear to have any significance in the selection of the viral genomes. Since its reporting, this mutation in the O type backbone was never encountered anymore in the sequences that became predominant henceforth, namely A2a and I/A3i. Instead, the S protein had acquired another mutation, L54F at a high frequency among A2a and A2 types where D614G is predominant. DynaMut analysis of L54F mutation has also identified it as a stabilizing one.

A2a subtype had acquired the greatest number of mutations in ORF1a compared to other subtypes. Among them S318L (nt C1218T) and Q676P (nt A2292C) in nsp2; S1515F (nt C4809T), I1159M (nt A3742G), S1534I (C4809T), A1812D (nt C5700A) and S2013F (nt C6573T) in nsp3; D3042N (nt G9389A) and A3143V (nt C9693T) in nsp4, S3517F (nt C10815T) in nsp5 and L3606F (nt G11083T) in nsp6 were highly frequent in the population. Three mutations in N gene S194L (nt C28854T), R203K (nt G28881A) and G204R (nt G28883C) were also found to be abundant within the A2a subtype. Furthermore, some researches indicated that a part of the nucleocapsid (N) protein of SARS-CoV (aa 161–211) is required for interacting with human cellular heterogeneous nuclear ribonucleoprotein A1 and this can play a regulatory role in the synthesis of SARS-CoV RNAs (Luo *et al.*, 2005). So, it would be interesting to see whether these mutations affect SARS-CoV-2 replication or not. Other mutations like L46F (nt C25528T) and Q57H (nt G25563T) in ORF3a were observed among the A2a isolates at high frequency. The 3a protein was predicted to be a transmembrane protein (Zeng *et al.*, 2004) and may be involved in ion channel formation during infection by co-localizing in the Golgi network (Lu, Xu and Sun, 2010). However, the Q57H mutation does not occur in the 6 defined domains of ORF3a (Issa *et al.*, 2020). It will be interesting to investigate whether these mutations have any role in virus transmission or replication.

From the perspective of the Indian isolates, occurrence of mutations in ORF1a was observed at higher number. Among the non-structural proteins of ORF1a, nsp3 (papain-like protease) tends to accumulate the highest number of mutations. When overall frequencies were compared, D614G in S protein and P323L in nsp12 were found to be highest along with a synonymous nucleotide change C241T in 5’UTR among all the Indian sequences (Fig 2).

SARS-CoV-2 genome is made up of 29.94% adenine, 18.37% guanine, 19.61% cytosine and 32.08% uracil. Compared to other RNA viruses (i.e. Dengue, Chikungunya) SARS-CoV-2 contains a higher amount of uracil (Fig 3). Interestingly, the most frequent changes in nucleotide were observed as C>T in case of non-synonymous mutations (Fig 4). This virus tends to change nucleotides into uracil at a high frequency which indicates the biasness of viral RdRp. These findings can help in selecting effective nucleoside/nucleotide analogues to test as effective antivirals. Analogously, herpes simplex virus (HSV) genome is GC-rich and 9-(2-hydroxyethoxymethyl) guanine (Acyclovir), the “gold-standard” herpes antiviral is incidentally a guanosine analogue. Acyclovir was the first highly virus-selective antiviral drug. It serves as a more preferential substrate for the HSV-encoded thymidine-kinase than host cell kinases for its initial phosphorylation (Frobert *et al.*, 2005; Jiao *et al.*, 2019). Previously, 3′-azido-2′,3′-unsaturated thymine analogue has shown better activity compared to other nucleoside analogues against SARS-CoV (EC50=10.3 µM) with a significant level of toxicity (Chu *et al.*, 2006). These findings will be helpful towards developing new antiviral candidates for SARS-CoV-2 where uracil/thymidine analogues may have an upper hand.

In summary, the identification and characterization of mutations in the SARS-CoV-2 genome will provide a better understanding of viral genome divergence and disease spread (Fig 5). The observations reported in this study require further experimental confirmation/validation.

**Fig 5:**
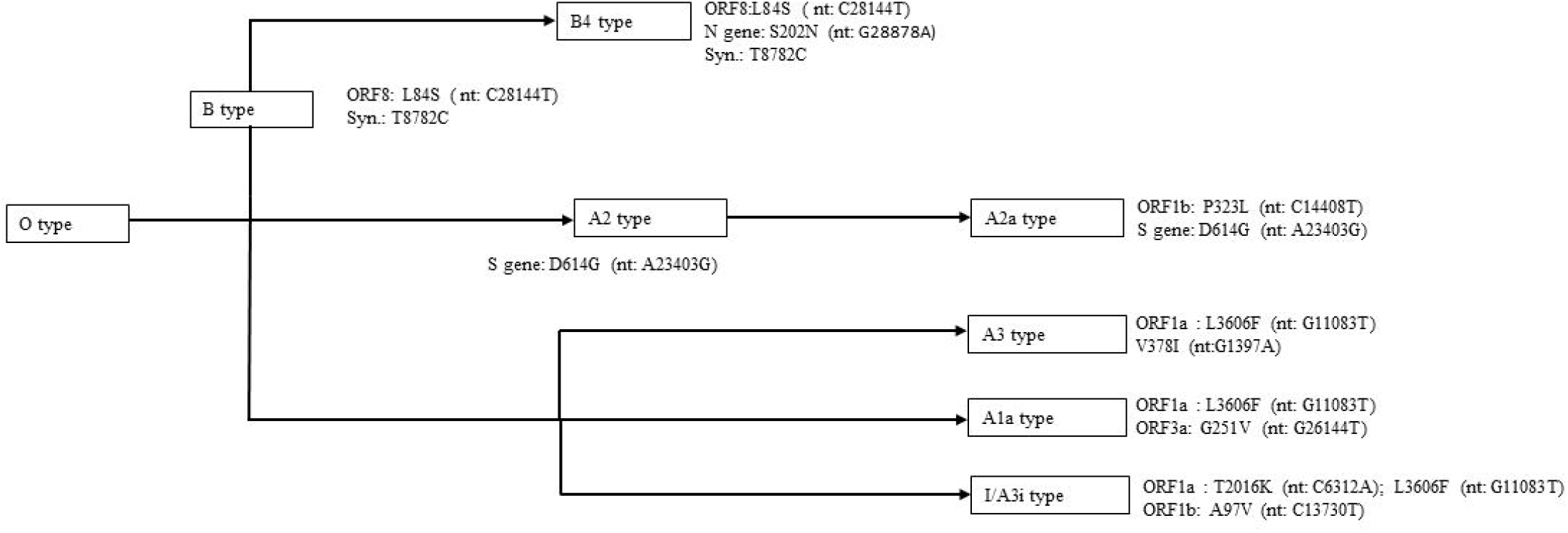
Schematic representation of SARS-CoV-2 types prevalent in the Indian population up to July 2020. It is based on the simplified understanding of the non-synonymous changes that shaped the emergence, divergence and prevalence of the different SARS-CoV-2 types.

## Credit authorship contribution statement

**Subrata Roy**- Sequence analysis, draft writing, visual representation

**Himadri Nath** & **Abinash Mallick**- DynaMut analysis, review and editing draft.

**Subhajit Biswas**- Conceptualization, Supervision and monitoring, Critical review and editing.

## Acknowledgement

SR acknowledges UGC for his Senior Research Fellowship. HN & AM thank CSIR for their SRF and JRF respectively. The authors acknowledge CSIR-IICB for providing laboratory facilities for conducting the present work.

## Funding

The project was funded by a grant from the **Council of Scientific and Industrial Research, India** to S.B. Grant number: MLP 130; CSIR Digital Surveillance Vertical for COVID-19 mitigation in India.

## Disclosure statement

The authors declare no competing interests.

## Supplementary data

**Supplementary Table 1. Indian SARS CoV 2 sequences and list of mutations**

## Notes

### Competing Interest Statement

The authors have declared no competing interest.

